## Supplement tables for "Seizure onset and offset pattern determine the entrainment of the cortex and substantia nigra in the nonhuman primate model of focal temporal lobe seizures"

Supplementary Table 1: Mean ± SEM values obtained in the pre-ictal, onset, offset and post-ictal periods in the HPC and SN for NHP1 and NHP 2. Statistical comparison performed with a Friedman repeated test and Dunnett’s for post hoc comparison with the values preceding the seizures, *<0.05, **<0.01, ***<0.001. Numbers in bold represent results consistent for both animals.

|  |  | NHP 1 | | | |  | NHP 2 | | | |
| --- | --- | --- | --- | --- | --- | --- | --- | --- | --- | --- |
|  |  | Pre-ictal | Onset | Offset | Post-ictal |  | Pre-ictal | Onset | Offset | Post-ictal |
| HPC | [1–7Hz] | 0.049±0.014 | **0.127±0.024 **** | 0.078±0.017 | 0.042±0.010 |  | 0.082±0.006 | **0.116±0.005***** | 0.069±0.004 | **0.049±0.002***** |
|  | [8–12Hz] | 0.010±0.001 | **0.060±0.008***** | **0.070±0.015***** | **0.038±0.008*** |  | 0.008±0.001 | **0.020±0.001***** | **0.018±0.001***** | 0.010±0.001 |
|  | [13–25Hz] | 0.003±0.001 | **0.030±0.003***** | **0.025±0.006**** | **0.054±0.012***** |  | 0.002±0.001 | **0.008±0.001***** | **0.005±0.001***** | 0.028±0.001 |
| SN | [1–7Hz] | 0.027±0.006 | 0.023±0.005 | 0.027±0.007 | 0.021±0.006 |  | 0.009±0.001 | 0.010±0.001 | 0.012±0.001 | 0.008±0.001 |
|  | [8–12Hz] | 0.004±0.001 | **0.006±0.001*** | 0.005±0.001 | 0.004±0.001 |  | 0.002±0.001 | **0.004±0.001***** | 0.003±0.001*** | 0.002±0.001 |
|  | [13–25Hz] | 0.001±0.001 | **0.003±0.001*** | 0.001±0.001 | 0.001±0.001 |  | 0.001±0.001 | **0.002±0.001***** | 0.001±0.001*** | 0.001±0.001 |

Supplementary Table 2­: Mean ± SEM values of the HPC/SN coherence obtained before, at the beginning and the end of the seizure for NHP1 and NHP 2. Statistical comparison performed with a Friedman repeated test and Dunnett’s for post hoc comparison with the pre-ictal values, *<0.05, **<0.01, ***<0.001. Numbers in bold represent results consistent for both animals.

|  |  |  | NHP 1 | | | |  | NHP 2 | | | |
| --- | --- | --- | --- | --- | --- | --- | --- | --- | --- | --- | --- |
| HPC-SN |  | Pre-ictal | | Onset | Offset | Post-ictal |  | Pre-ictal | Onset | Offset | Post-ictal |
|  | [1–7Hz] | 0.55±0.05 | | 0.56±0.04 | 0.52±0.03 | 0.53±0.03 |  | 0.60±0.01 | 0.55±0.01* | 0.51±0.01*** | 0.53±0.01* |
|  | [8–12Hz] | 0.57±0.04 | | 0.57±0.03 | 0.55±0.03 | 0.60±0.03 |  | 0.53±0.01 | 0.48±0.01 | 0.47±0.01 | 0.50±0.01 |
|  | [13–25Hz] | 0.50±0.03 | | **0.55±0.02*** | 0.54±0.01* | 0.60±0.03* |  | 0.53±0.01 | **0.62±0.01***** | 0.53±0.01** | 0.50±0.01** |

Supplementary Table 3: Mean ± SEM spectra values and coherence obtained in the pre-ictal and onset periods in the HPC and SN for LAF and HAS onset patterns. Statistical comparison performed with a Wilcoxon Signed Rank test for paired quantitative data, *<0.05, **<0.01, ***<0.001. Comparisons between LAF and HAS seizures were performed with a Mann-Whitney Rank Sum test (# <0.05). Statistical values were corrected for multiple comparison using Bonferroni correction.

|  |  | LAF (n=44) | |  |  | HAS (n=18) | |
| --- | --- | --- | --- | --- | --- | --- | --- |
|  |  | Pre-ictal | Onset |  |  | Pre-ictal | Onset |
| HPC | [1–7Hz] | 0.063±0.006 | 0.107±0.006 *** |  |  | 0.104±0.014# | 0.137±0.009**# |
|  | [8–12Hz] | 0.007±0.001 | 0.029±0.004*** |  |  | 0.010±0.001 | 0.023±0.002*** |
|  | [13–25] | 0.002±0.001 | 0.013±0.002*** |  |  | 0.003±0.001 | 0.009±0.001*** |
| SN | [1–7Hz] | 0.010±0.001 | 0.011±0.001 |  |  | 0.015±0.003 | 0.016±0.003 |
|  | [8–12Hz] | 0.002±0.001 | 0.004±0.001*** |  |  | 0.003±0.001 | 0.005±0.001* |
|  | [13–25Hz] | 0.001±0.001 | 0.002±0.001*** |  |  | 0.001±0.001 | 0.004±0.001***# |
| HPC/SN | [1–7Hz] | 0.56±0.02 | 0.54±0.01 |  |  | 0.68±0.03# | 0.59±0.02** |
|  | [8–12Hz] | 0.52±0.01 | 0.50±0.01 |  |  | 0.58±0.03 | 0.50±0.03 |
|  | [13–25Hz] | 0.55±0.01 | 0.59±0.01* |  |  | 0.63±0.02# | 0.63±0.02 |

Supplementary Table 4: Mean ± SEM spectra values and coherence obtained in the offset and post-ictal periods in the HPC and SN for ARR, RHY and BS offset patterns. Statistical comparison performed with a Wilcoxon Signed Rank test for paired quantitative data, *<0.05, **<0.01, ***<0.001.

|  |  | ARR (n=36) | |  | RHY (n=44) | |  | BS (n=12) | |
| --- | --- | --- | --- | --- | --- | --- | --- | --- | --- |
|  |  | Offset | Post-ictal |  | Offset | Post-ictal |  | Offset | Post-ictal |
| **HPC** | [1–7Hz] | 0.067±0.004 | 0.047±0.003*** |  | 0.079±0.011 | 0.047±0.006*** |  | 0.056±0.006 | 0.047±0.007 |
|  | [8–12Hz] | 0.021±0.002 | 0.011±0.002*** |  | 0.046±0.011 | 0.025±0.006*** |  | 0.015±0.001 | 0.009±0.002*** |
|  | [13–25] | 0.006±0.002 | 0.003±0.001*** |  | 0.017±0.004 | 0.034±0.009** |  | 0.005±0.001 | 0.002±0.001*** |
| **SN** | [1–7Hz] | 0.011±0.002 | 0.007±0.001*** |  | 0.014±0.002 | 0.011±0.002* |  | 0.017±0.002 | 0.013±0.001* |
|  | [8–12Hz] | 0.003±0.001 | 0.002±0.001*** |  | 0.004±0.001 | 0.003±0.001*** |  | 0.004±0.001 | 0.003±0.001*** |
|  | [13–25] | 0.0006±0.002 | 0.0003±0.001*** |  | 0.001±0.001 | 0.0006±0.001*** |  | 0.0008±0.001 | 0.0005±0.003*** |
| **HPC/SN** | [1–7Hz] | 0.53±0.014 | 0.53±0.012 |  | 0.53±0.020 | 0.55±0.019 |  | 0.46±0.011 | 0.56±0.019*** |
|  | [8–12Hz] | 0.47±0.011 | 0.52±0.013** |  | 0.53±0.021 | 0.55±0.024 |  | 0.050±0.02 | 0.048±0.015 |
|  | [13–25] | 0.51±0.012 | 0.53±0.008 |  | 0.54±0.014 | 0.54±0.011 |  | 0.46±0.016 | 0.53±0.008*** |

Supplementary Table 5: Mean ± SEM values obtained before, at the beginning and the end of the seizures in the SI for LAF and HAS onset patterns. Statistical comparison performed with a Friedman repeated test and Dunnett’s for post hoc comparison with the values preceding the seizures, *<0.05, **<0.01, ***<0.001. Comparisons between LAF and HAS seizures were performed with a Mann-Whitney Rank Sum test (# <0.05). Statistical values were corrected for multiple comparison using Bonferroni correction.

|  |  | LAF (n=36) | |  |  | HAS (n=15) | |
| --- | --- | --- | --- | --- | --- | --- | --- |
|  |  | Pre-ictal | Onset |  |  | Pre-ictal | Onset |
| SI | [1–7Hz] | 0.024±0.002 | 0.033±0.004 ** |  |  | 0.012±0.002### | 0.016±0.004## |
|  | [8–12Hz] | 0.021±0.003 | 0.026±0.003 |  |  | 0.012±0.002# | 0.012±0.003## |
|  | [13–25] | 0.004±0.001 | 0.005±0.001* |  |  | 0.002±0.001## | 0.003±0.001## |
| HPC/SI | [1–7Hz] | 0.46±0.01 | 0.53±0.01*** |  |  | 0.46±0.02 | 0.52±0.02* |
|  | [8–12Hz] | 0.44±0.01 | 0.43±0.01 |  |  | 0.43±0.02 | 0.41±0.01 |
|  | [13–25Hz] | 0.44±0.01 | 0.43±0.01 |  |  | 0.43±0.02 | 0.42±0.01 |

Supplementary Table 6: Mean ± SEM values obtained before, at the beginning and the end of the seizures in the SI for LAF and HAS onset patterns. Statistical comparison performed with a Friedman repeated test and Dunnett’s for post hoc comparison with the values preceding the seizures, *<0.05, **<0.01, ***<0.001. Comparisons between LAF and HAS seizures were performed with a Mann-Whitney Rank Sum test (# <0.05). Statistical values were corrected for multiple comparison using Bonferroni correction.

|  |  | ARR (n=35) | |  | RHY (n=9) | |  | BS (n=12) | | |
| --- | --- | --- | --- | --- | --- | --- | --- | --- | --- | --- |
|  |  | Offset | Post-ictal |  | Offset | Post-ictal |  | Offset | Post-ictal | |
| SI | [1–7Hz] | 0.012±0.002 | 0.011±0.002 |  | 0.033±0.007 | 0.028±0.005 |  | 0.022±0.004 | | 0.021±0.003 |
|  | [8–12Hz] | 0.013±0.002 | 0.010±0.001 |  | 0.026±0.004 | 0.020±0.002 |  | 0.024±0.003 | | 0.018±0.003 |
|  | [13–25] | 0.002±0.001 | 0.002±0.001 |  | 0.005±0.001 | 0.004±0.001 |  | 0.004±0.001 | | 0.004±0.001 |
| HPC/SI | [1–7Hz] | 0.44±0.01 | 0.46±0.01 |  | 0.56±0.03 | 0.53±0.03 |  | 0.48±0.02 | | 0.47±0.02 |
|  | [8–12Hz] | 0.41±0.01 | 0.43±0.01 |  | 0.44±0.01 | 0.47±0.01 |  | 0.43±0.01 | | 0.45±0.01 |
|  | [13–25Hz] | 0.42±0.01 | 0.43±0.01 |  | 0.44±0.01 | 0.47±0.01 |  | 0.42±0.01 | | 0.44±0.01 |
